# A comprehensive EEG dataset for investigating visual touch perception

**DOI:** 10.1101/2025.07.10.664069

**Authors:** Sophie Smit, Almudena Ramírez-Haro, Manuel Varlet, Denise Moerel, Genevieve L. Quek, Tijl Grootswagers

**Affiliations:** The MARCS Institute for Brain, Behaviour and Development, Western Sydney University, Australia; School of Psychology, Western Sydney University, Australia; School of Computer, Data and Mathematical Sciences, Western Sydney University, Australia

## Abstract

Visual touch perception is fundamental to body awareness and social bonding, helping us perceive ourselves and connect with others. However, understanding its neural basis has been limited by a lack of comprehensive public datasets. To address this, we present an electroencephalography (EEG) and self-report dataset from 80 participants. It includes neural responses from 2,880 trials per participant, during which participants viewed touch interactions between two hands involving either direct skin contact or object-mediated touch. Complementing the EEG data are self-report measures on empathy, perspective-taking, vicarious touch, and mirror-touch synaesthesia, providing a robust resource for exploring individual differences in visual touch perception. Ninety unique videos were presented in four orientations, with trials evenly split between self- and other-perspectives, and showing touch to either the left or right hand. This design supports investigation into how viewing perspective and hand laterality influence touch processing. Technical validation reveals distinct neural patterns associated with visual, sensory, and emotional touch dimensions, highlighting the dataset’s potential to advance research into diverse aspects of visual touch perception.

## Background & Summary

In cognitive neuroscience, interest in the perception of touch through visual cues has grown, particularly in relation to empathy for others’ sensations ^1–4^, touch communication ^5,6^, and multisensory body perception ^7–9^. With the rise of online interactions ^10,11^, research has further expanded to examine whether visually perceiving touch can replicate some benefits of physical touch ^12–17^. This focus on understanding the nuances of visual touch processing is thus increasingly relevant both for fundamental science and practical applications. Advancing our knowledge of how the brain processes observed touch requires comprehensive, publicly available datasets. While neuroimaging datasets are increasingly accessible^18^, few target human touch perception specifically. Prior studies have often relied on static or simplified visual touch stimuli to control experimental variables ^19–26^, limiting their reflection of dynamic real-world touch interactions. Moreover, many studies have used small sample sizes (< 20 participants) and have restricted data access, underscoring the need for a high-quality, open-access dataset that captures the complexity of visual touch perception.

We present a comprehensive 64-channel electroencephalography (EEG) dataset from 80 participants who viewed dynamic videos of close-up touch interactions between two hands, taken from the Validated Touch-Video Database ^27^. Visual elements in these videos (e.g., hand, background) were kept consistent to isolate sensory and emotional aspects of touch processing. Touch either involved direct skin contact or showed object-mediated touch. These videos were systematically rated in a prior study by an independent sample on dimensions such as valence, arousal, and threat ^27^. For this dataset, we also manually coded the videos for 12 distinct touch types (e.g., touch, stroke, scratch), 28 objects (e.g., brush, knife, hammer), and 8 materials (e.g., cotton, plastic, metal). Some videos depicted one hand approaching the other, while others showed contact from the start. Using a rapid sequence design ^28,29^, participants viewed a total of 90 unique videos. Each video was presented in four orientations (achieved by horizontal and/or vertical flipping to depict a left or right hand and a self-or other-perspective), creating 360 distinct stimuli. Each of these 360 stimuli was repeated eight times, resulting in a total of 2,880 such trials over a 55-minute session. Participants were instructed to count additional target videos (depicting touch to a white block) to ensure engagement throughout the task. Each video lasted 600 ms, followed by a 200 ms inter-stimulus interval to allow complete neural responses without forward or backward masking ^30^. Technical validation using multivariate pattern analysis ^31,32^ revealed distinct neural patterns associated with self-versus other-perspectives, sensory properties, and emotional content of the observed touch, at both the group and individual levels. Additionally, we collected self-report measures assessing empathy, perspective-taking, vicarious touch, and mirror-touch synaesthesia—factors previously studied in visual touch processing ^1,21,24,33,34^. This dataset, combining neural and self-report data, provides a robust foundation for advancing research into the mechanisms of visual touch perception.

This dataset provides a comprehensive resource for exploring the complexities of human visual touch perception. It contains neural responses to a wide range of observed naturalistic touch interactions. This diversity allows researchers to investigate how different types of touch are processed in the brain, including distinctions between positive and negative touch experiences and the effects of different touch mediums. The stimuli are presented from both self- and other- perspectives, enabling investigations into how perspective-taking influences touch perception. With data from a large number of participants and trials, this high-powered dataset supports robust analyses focused on specific research areas such as empathy, sensory processing, and emotional responses to observed touch. The combination of EEG recordings and self-report measures allows for nuanced exploration of individual differences in how observed touch is processed. We expect that this publicly available dataset will enable collaboration, replication, and the development of new insights.

## Methods

A total of 80 individuals participated in the experiment, either in exchange for course credit or a payment of AUD$45. The final sample included 54 women, 24 men, and 2 non-binary, with a mean age of 30.08 years (SD = 10.77) and an age range of 18-76 years. Handedness distribution was as follows: 68 right-handed, 9 left-handed, and 3 ambidextrous. All participants reported normal or corrected-to-normal vision. The study received ethical approval from the Western Sydney University Human Research Ethics Committee (project number: H15644), and informed written consent was obtained from all participants at the start of the experiment.

The stimuli for this experiment were sourced from the Validated Touch-Video Database ^27^, which comprises 90 videos of tactile interactions between two hands, filmed from a first-person perspective. Each video shows a right hand contacting a left hand, either directly or using an object, and varies across dimensions such as the type of touch (e.g., stroking, pressing, stabbing), the use of objects (e.g., brush, hammer, knife), and the material involved (e.g., cotton, plastic, metal). The database has been previously validated by a large sample of participants ^27^, who categorised each video based on hedonic qualities (neutral, pleasant, unpleasant, painful) and rated levels of arousal and perceived threat. We also manually coded each video for touch type (12), object (28), and material (8). The original videos varied in length; for the purpose of this dataset, all videos were trimmed to 600 ms to ensure consistency for EEG analysis, with the touch event centred within the clip. To introduce variations in perspective, the videos were presented in four orientations, created by applying horizontal, vertical, or combined flips. This resulted in a total of 360 unique stimuli that depicted touch directed toward either the left or right hand from both a self-or other-perspective. We previously confirmed that modifications to the videos, including shorter duration, smaller presentation size, and different orientations, did not affect the rated attributes of the touch compared to the original versions (mean across modifications: *r* = 0.89, see ^35^). In this study, we used the ratings for the original videos from the Validated Touch-Video Database ^27^, as these have been validated by a large sample. Both the adapted videos used here, and original videos and validation data, are available online (https://osf.io/jvkqa/). Each video consisted of 15 frames, presented at 25 frames per second on a 120 Hz display. The resolution was set to 256×144 pixels, and the stimuli were viewed from a distance of approximately 60 cm, corresponding to a visual angle of 6.2°.

We employed a rapid sequence design ^28,29^ where each participant viewed 32 sequences of 90 videos selected from the 360 unique stimuli (Fig. 1), presented in a fully randomised order. Each video was repeated eight times per participant, resulting in 2,880 trials in total. To ensure participants remained attentive, additional target stimuli—videos featuring touch interactions between a hand and a white object rather than another hand—were randomly embedded among the main touch videos. Each sequence contained between 1 and 9 target stimuli, with at least 12 non-target videos separating target trials. Participants were instructed to count the target videos and report their total at the end of each sequence using the top row of the keyboard, after which they received immediate feedback on their accuracy. On average, participants achieved 81.21% accuracy (SD = 18.83%, min-max = 6.25% - 100%) in the target detection task, reflecting high engagement throughout the experiment. Each video was separated by a 200 ms gap, and sequence durations varied from 54 to 78 seconds, depending on the number of target stimuli included. Participants viewed the stimuli on a 24-inch ViewPixx monitor against a grey background (Fig. 1B) and were encouraged to take breaks between sequences and resumed the task by pressing a key. The whole task lasted approximately 55 minutes. The experiment was programmed and run using Python and PsychoPy version 2023.3.1 ^36^. EEG data were continuously recorded using a 64-channel BioSemi Active-Two system at a sampling rate of 2048 Hz (BioSemi, Amsterdam, The Netherlands). Voltage offsets were maintained at ±20 mV, and electrodes were positioned according to the international 10/20 system ^37,38^.

**Figure 1.**
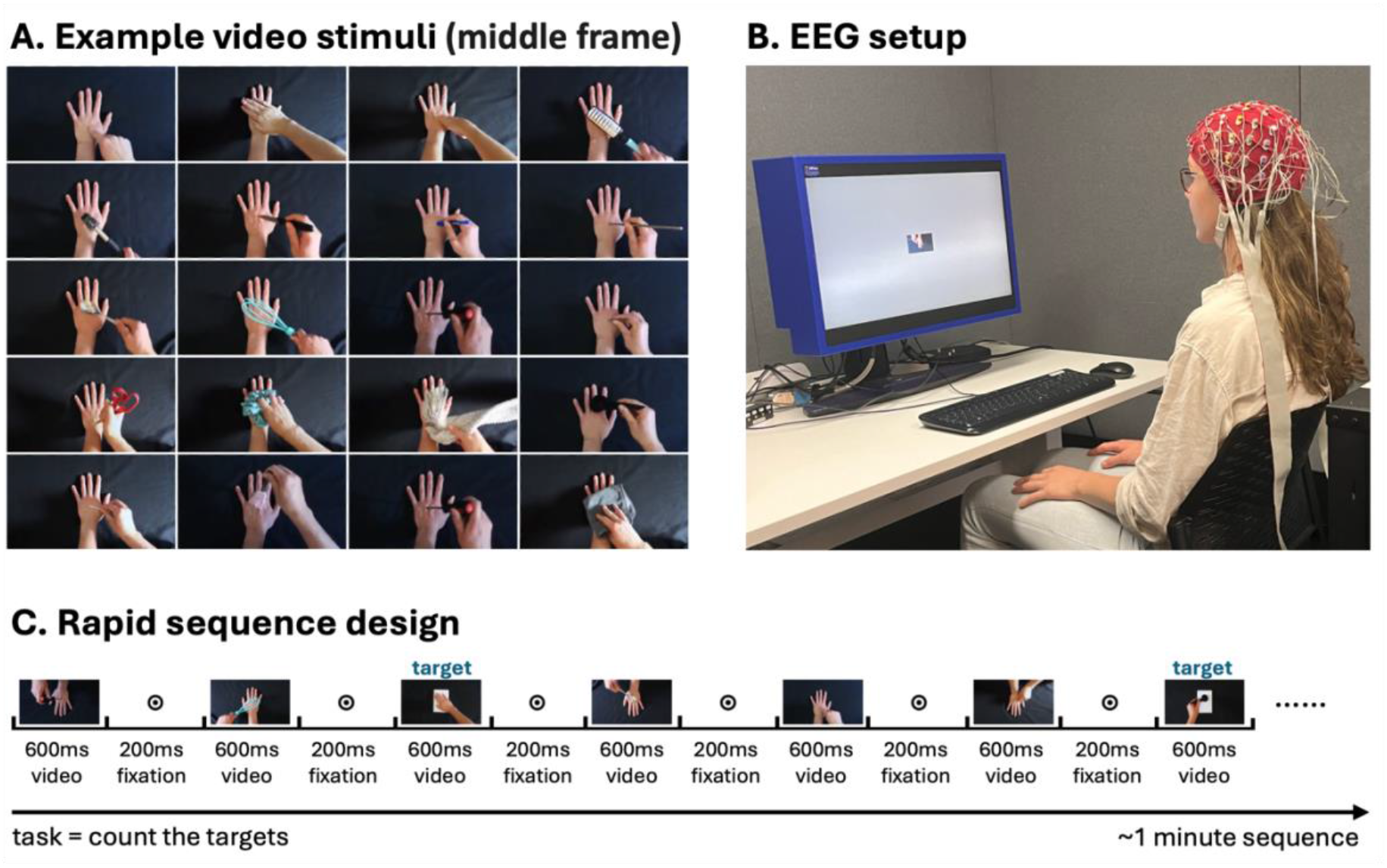
Example stimuli, EEG setup, and experimental design. (A) Middle frame of a selected subset of touch videos used in the experiment, depicting the range of tactile interactions. (B) EEG experimental setup (image depicts one of the authors). (C) Rapid sequence design, illustrating a portion of the sequence in which visual stimuli are rapidly presented during the experiment.

To complement the neural data, we included four self-report questionnaires that assess individual differences in touch perception and other traits, administered in the order described here. These took an additional 15-30 minutes to complete and were administered during EEG setup (i.e., prior to EEG recording). An overview of all the questions can be found on the Open Science Framework (OSF) repository for this project. 1) The **Vicarious Touch Questionnaire** ^24,34^ asks participants if they felt sensations while watching 10 videos of neutral touch to a hand from the Validated Touch-Video Database ^27^, including the type of sensation, the intensity, the hedonic quality, and where on the body it was felt. 2) The **Short Empathy Quotient (EQ) Scale** ^39^ measures empathy through three five-question subscales: cognitive empathy (e.g., “I am good at predicting how someone will feel”), social skills (e.g., “I find it hard to know what to do in a social situation”), and emotional reactivity (e.g., “Seeing people cry does not really upset me”). 3) The **Interpersonal Reactivity Index (IRI) Perspective-Taking Subscale** ^40^ assesses participants’ ability to adopt others’ viewpoints via seven questions such as “I believe that there are two sides to every question and try to look at them both.” 4) A **Short Mirror-Touch Synaesthesia Questionnaire** was developed for this study to identify individuals who experience mirror-touch synaesthesia, where observing touch on someone else reliably elicits a similar sensation in the observer ^33^. The questionnaire explores whether participants think they have this condition, and if so, the nature of their experiences, the frequency with which vicarious touch occurs, and the location on the body where the touch is felt. These self-report measures were included to explore how individual differences in empathy, perspective-taking, and vicarious touch experiences relate to variations in neural processing.

### Data Records

Raw and preprocessed EEG data, self-report questionnaire responses, and individual decoding results are available on OpenNeuro in BIDS format (https://openneuro.org/datasets/ds005662). Custom code for generating the figures in this paper, along with an overview of all questionnaires and the files needed to run the experiment, is provided on OSF (https://osf.io/t6nfa/). The video stimuli used here were adapted from the Validated Touch-Video Database ^27^, alongside validation data for the original videos (see Methods). Both stimulus sets are available on the OSF page of the Validated Touch-Video Database (https://osf.io/jvkqa/) ^27^.

### Technical Validation

We performed basic quality checks and technical validation on both the individual- and group-level data. Epochs were extracted from -100 to 800 ms relative to stimulus onset, downsampled to 200 Hz, and baseline-corrected using the -100 to 0 ms window. We applied a 1-100 Hz band-pass filter, re-referenced the data to the common average, and smoothed the time series using a moving average filter over 10 samples (equivalent to 40 ms). All voltage readings from each channel and time point were retained for analysis, and no additional preprocessing steps were performed for the technical validation analyses presented here. All preprocessing and subsequent analyses were conducted using the MNE-Python toolbox version 1.7 ^31^.

We applied standard time-series multivariate pattern analysis (MVPA) ^31,32^ to decode information present in neural activation patterns. Decoding analyses were first performed separately for each participant, and decoding accuracies were then averaged across participants to assess group-level patterns. Specifically, we decoded three dimensions: viewing perspective (self vs. other), the material of the object used for touch (8 categories; see Methods for details), and valence (continuous rating). For valence, we used normative ratings from the Validated Touch-Video Database for the original videos ^27^, where an independent sample of participants categorised each video as pleasant or unpleasant. We performed a Principal Component Analysis (PCA) on the proportion of participants who classified each video as pleasant or unpleasant; the first principal component was extracted to derive a continuous composite valence score. In addition, we decoded hand orientation (left vs. right hand; self-vs. other-perspective) to explore variability at the individual level. Classification analyses were used for categorical variables (perspective, material, and orientation), and regression for the continuous valence scores (see also ^35^). This decoding approach enabled us to assess whether and how different types of information are represented in neural patterns across individuals and, by averaging accuracies, at the group level.

Group-mean decoding accuracies (Fig. 2, top panel) reveal that information about perspective (self vs. other) peaks around 150 ms, while information about the material of the touching object and valence peak around 300 ms, indicating that body cues, sensory, and emotional aspects of touch are encoded in the neural data. Figure 2 (bottom panel) shows individual variance in decoding accuracies. Similarly, at the individual-subject level (Fig. 3), hand orientation (left vs. right hand; self-vs. other-perspective) was decoded above chance for every participant at some point in the trial. While most participants showed peak decoding around 135 ms (group-mean accuracy at that time point: 34%; chance level = 25%) the plot highlights some variability in the timing of above-chance decoding across participants. These findings demonstrate the encoding of body cues, sensory properties, and affective dimensions in neural patterns, highlighting the EEG database’s potential for examining various facets of touch perception at both individual and group levels.

**Figure 2.**
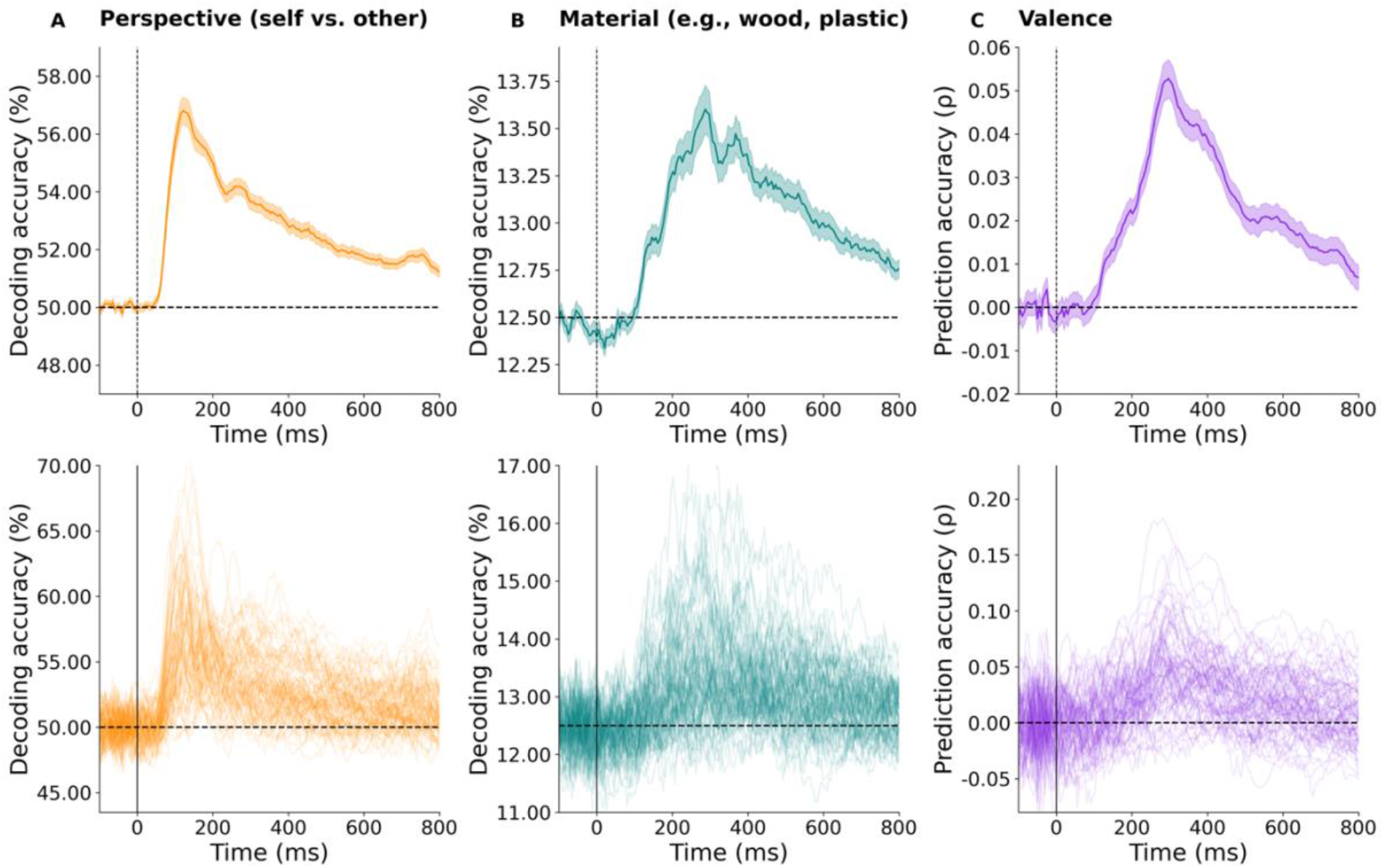
Group-level EEG dynamics and decoding analysis. (A) Decoding of hand perspective (self vs. other), (B) object material, and (C) valence. The top row displays group-averaged decoding accuracies over time, with shaded areas indicating the standard error of the mean, while the bottom row shows decoding accuracies for individual participants. Dashed horizontal lines indicate chance levels and vertical lines mark stimulus onset at 0 ms.

**Figure 3.**
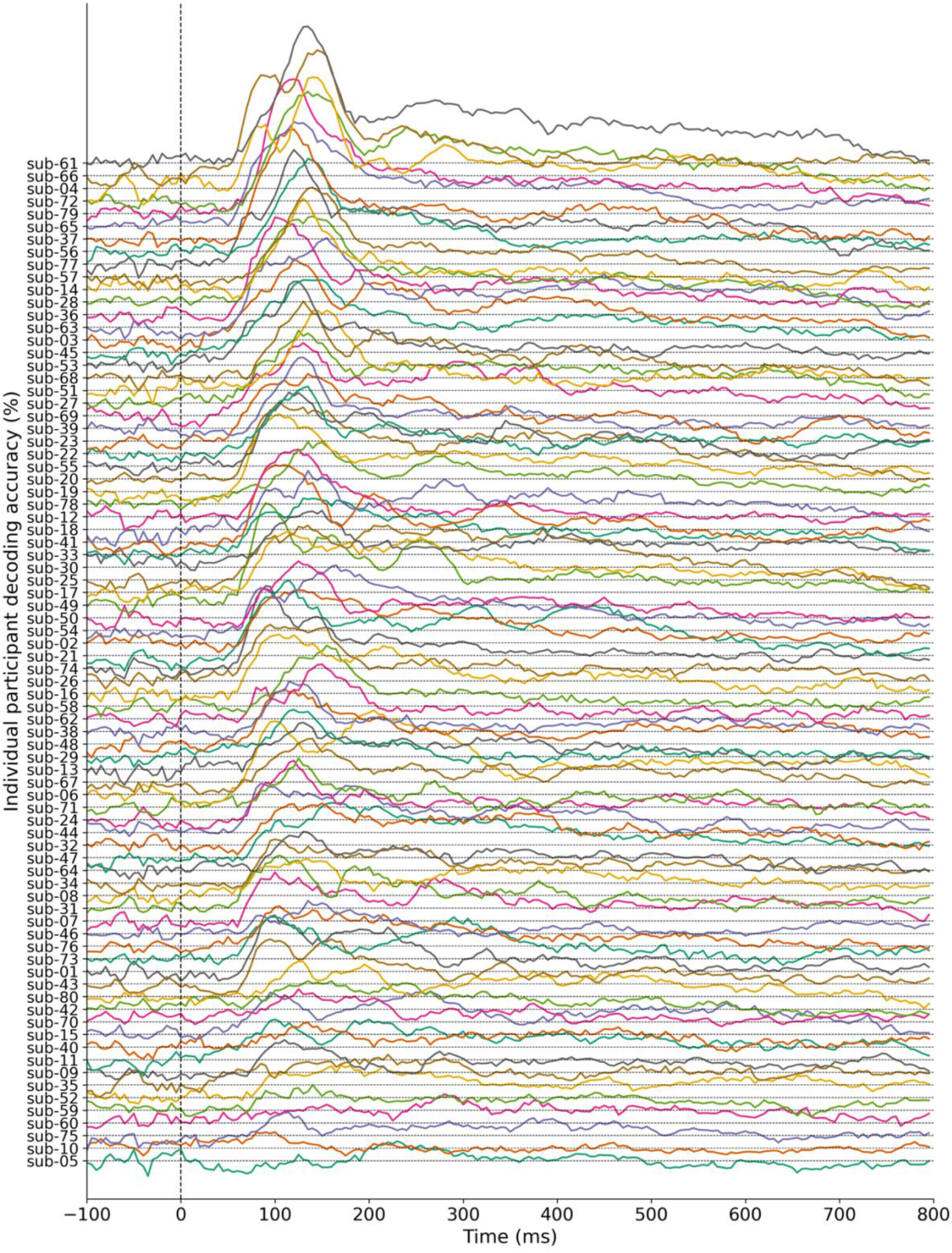
Participant-level decoding accuracies for four hand orientations. Each line represents a participant’s decoding accuracy for hand orientation (4 options: left vs. right hand; self-vs. other-perspective) over time, with distinct colours differentiating participants. Participants are sorted based on their individual decoding accuracy at the group-level peak time point of 135 ms. This allows visual comparison of decoding timing variability across individuals. Dashed horizontal lines indicate chance levels and the vertical line mark stimulus onset at 0 ms.

Self-report data reveal a wide range of responses across participants. Scores from both the EQ and IRI questionnaires demonstrate a well-distributed range of empathy and perspective-taking levels in our sample (Fig. 4A, B). Of all participants, 41.3% reported experiencing a vicarious sensation in their own hand on at least 1/10 trials while observing touch to a hand in 10 different videos (Fig. 4C). This result closely matches previous findings using the same methods, which reported a 47.1% prevalence of vicarious touch in a typical sample ^24^. Additionally, 21.2% of participants indicated that they might have mirror-touch synaesthesia (Fig. 4D), which is higher than the typical ∼10% prevalence of self-reported cases ^33,34,41^. However, follow-up responses suggested that many of these participants described general, non-localised sensations throughout the body, which is inconsistent with classic mirror-touch synaesthesia ^33^. Adjusting for this distinction, 10% of participants in our sample reported both a belief that they may have the condition and that they mostly experience localised tactile sensations when observing touch. Overall, these findings suggest that this dataset is well-suited for exploring individual differences in touch perception and related traits.

**Figure 4.**
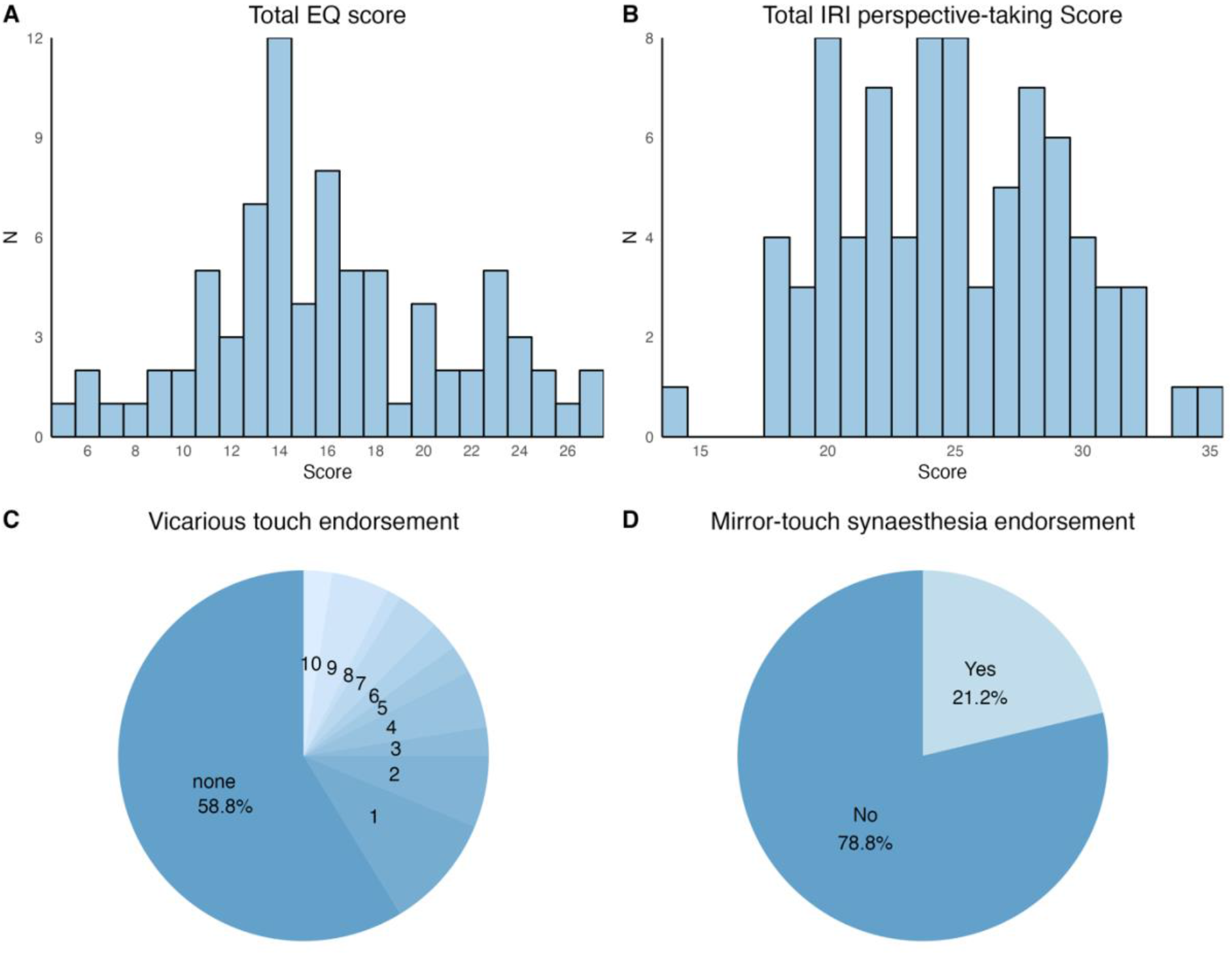
High-level summary of self-report results. (A) Histogram of total Empathy Quotient (EQ) scores across participants, summing points from 15 items on the Cognitive, Social Skills, and Emotional Reactivity subscales. Responses were scored as 0, 0, 1, or 2, with “strongly disagree” and “somewhat disagree” receiving 0 points, “somewhat agree” 1 point, and “strongly agree” 2 points (or reversed for negatively phrased items, following the original scoring). The maximum score is 30, with higher scores reflecting greater empathy. (B) Histogram of Interpersonal Reactivity Index (IRI) Perspective-Taking subscale scores. Participants responded to 7 items with scores ranging from 0: “Does not describe me well” to 5: “Describes me very well”. The maximum score is 35 with higher scores reflecting greater perspective-taking ability. (C) Pie chart showing the distribution of trials (none–10) where participants reported vicarious touch on one or both hands. (D) Pie chart showing the percentage of participants who answered “yes” to the question: “Do you think you have mirror-touch synaesthesia?”. Participants who responded “yes” were asked follow-up questions about the nature of the experience, the context in which it occurs, and the location of the touch on the body.

## Code availability

Code and instructions to reproduce the technical validation analyses and figures presented in this manuscript are available on OSF (https://osf.io/t6nfa/), with all data available on OpenNeuro (https://openneuro.org/datasets/ds005662) as described above.

## Acknowledgements

This research was supported by Australian Research Council grants DP220103047 (MV) and DE230100380 (TG).

## Author Contributions

Conceptualisation: S.S., T.G.: Stimulus and task development: S.S., T.G.; Data acquisition: S.S., A.R.H.; Formal analysis: S.S., T.G.; Writing (original draft): S.S.; Writing (reviewing and editing): all authors.

## Competing interests

The authors declare no competing interests.

## Notes

https://osf.io/t6nfa/

https://openneuro.org/datasets/ds005662

https://osf.io/jvkqa/

### Competing Interest Statement

The authors have declared no competing interest.

